# Diphenhydramine Disrupts Sleep Architecture in 5XFAD Alzheimer’s Disease Model and Wild-Type Mice

**DOI:** 10.64898/2026.07.22.739929

**Authors:** Margaret H. Copeland, Diane E. Youngstrom, Kathryn S. Konrad, Graham H. Diering, Ayland C. Letsinger, Leslie R. Aksu, Jerrel L. Yakel, Jesse D. Cushman

## Abstract

Sleep disruption is common in Alzheimer’s disease (AD). Diphenhydramine (DPH), a first-generation antihistamine with anticholinergic properties, is widely used as an over-the-counter sleep aid. We tested whether chronic DPH treatment alters sleep architecture in 5XFAD and wild-type mice. Female 5XFAD (n=16) and wild-type (WT) littermates (n=14) were implanted with wireless telemetry recording devices to measure electroencephalography (EEG), electromyography (EMG), temperature, and activity continuously. After a 24h baseline recording at 5 months of age, mice received oral DPH (10 mg/kg) or vehicle at ZT0 for one month. After this chronic treatment, sleep was recorded continuously for 48h during ongoing dosing. Sleep was scored as rapid eye movement (REM), non-rapid eye movement (NREM) 1, NREM2, or wake. A survival curve analysis was used to investigate the microarchitecture of sleep phases after chronic diphenhydramine treatment. At baseline, 5XFAD mice had more time in NREM1 than WT controls and had shorter REM and NREM2 bouts. Chronic DPH treatment fragmented NREM2 in both genotypes, reducing long NREM2 bouts. DPH increased total duration of NREM1 and REM during the active phase, which is analogous to daytime drowsiness in humans. DPH did not rescue 5XFAD sleep deficits; instead, DPH treatment exacerbated NREM2 fragmentation. Overall, chronic DPH use degrades sleep quality and increases fragmentation in both WT and AD-model mice, which questions the use of sedating anticholinergics as sleep aids, especially in AD.

## 1. Introduction

Alzheimer’s disease (AD) is a progressive neurodegenerative disease characterized by cognitive decline, memory impairment, and widespread accumulation of pathological protein aggregates such as amyloid-beta (Aβ) plaques and tau neurofibrillary tangles. In addition to these canonical pathological features, sleep disturbances are increasingly recognized as a major component of AD progression. Individuals with AD frequently experience fragmented sleep, insomnia, excessive daytime sleepiness, and disruptions in circadian rhythms, contributing to cognitive decline and reducing quality of life (Himali et al., 2023; Zhang et al., 2022; Ju et al., 2014; Wang et al., 2020, Hita-Yañez et al., 2012; Hita-Yañez et al., 2013). Recent data from the Mayo Clinic suggest that 25% of individuals with mild to moderate dementia and nearly 50% of those with severe dementia experience sleep disruptions, further underscoring the pervasiveness of this issue (Mayo Clinic, 2024). Sleep disturbances precede cognitive impairment in many AD patients (Atayde et al., 2022; Gonzales et al., 2024).

Beyond being a symptom of AD, mounting evidence suggests that sleep disturbances may actively contribute to disease progression. Studies indicate that Aβ accumulation in the brain disrupts sleep architecture, reducing sleep efficiency and increasing wakefulness (Huang et al., 2012; Roh et al., 2012). The reverse is also true; sleep deprivation or chronic sleep disruption exacerbates Aβ aggregation and clearance deficits, forming a vicious cycle wherein disrupted sleep accelerates AD pathology, which in turn further impairs sleep (Lacerda et al., 2024; Ju et al., 2014; Wang et al., 2020; Tabuchi et al., 2015; Dissel et al., 2017). This vicious cycle can be thwarted by sleep aids. For example, orexin antagonists can enhance nonrapid eye movement (NREM) sleep leading to neuroprotective effects in tauopathy model mice (Parhizkar et al., 2025). However, not all sleep aids slow AD pathology. Benzodiazepines and sedating first-generation antihistamines may promote AD-like brain atrophy and even, paradoxically, decrease sleep quality (Barbaux et al., 2025; de Mendonça et al., 2023; Legrand et al., 2025; Risacher et al., 2016; Gray et al., 2015; Boyle et al., 2006; Katayose et al., 2012; Church et al., 2010). Given the close relationship between sleep and AD, it is critical to understand the efficacy of popular sleep aids and their effects on AD pathology.

There is growing concern about chronic use of diphenhydramine hydrochloride (DPH) as a sleep aid, especially in patients with Alzheimer’s disease (Kan et al., 2025; Risacher et al., 2016; Gray et al., 2015; Katayose et al., 2012; Church et al., 2010). DPH is a widely available over-the-counter H1-antihistamine. In addition to blocking histamine receptors, DPH antagonizes muscarinic acetylcholine (ACh) receptors. Because DPH blocks these receptors and crosses the blood-brain barrier, it has sedative properties and is commonly used as a sleep aid. Paradoxically, growing evidence suggests DPH may have no effect or even a harmful effect on sleep quality. For example, DPH does not significantly improve sleep latency or total sleep time compared to placebo, so it is not recommended for treatment of insomnia (Sateia et al, 2017). Also, first-generation antihistamines may suppress REM sleep, increase REM latency, and suppress cognitive performance the next day (Katayose et al., 2012; Church et al., 2010; Boyle et al., 2006, Rojas-Zamorano et al., 2009). Most concerningly, long-term use of strong anticholinergics, and DPH in particular, is linked with dementia incidence (Gray et al., 2015, Risacher et al., 2016).

Though evidence against DPH as a sleep aid is accumulating, many findings implicating DPH in pathology are based on epidemiological data (Gray et al., 2015, Risacher et al., 2016), so a causal effect has not been established between DPH and sleep impairment. Here, we sought to investigate the impact of chronic DPH use on sleep quality using the 5XFAD AD mouse model.

By using this model, we can establish the causal effects of DPH on sleep, which can clarify correlational epidemiological findings. This method also allows us to investigate more granular sleep changes caused by DPH through extended electroencephalogram (EEG) paired with electromyogram (EMG) recordings, which report sleep states in rodents to a high degree of precision. Females were used exclusively in this study as woman are twice as likely to be affected by AD as males (Alzheimer’s Association, 2017). Female 5XFAD mice mirror this pathology as they show more profound pathology than males; they exhibit higher levels of amyloid-beta plaque deposition, enhanced astrogliosis, and higher expression of 5XFAD transgenes (Devi et al., 2010; Poon et al., 2023; Sadleir et al., 2015; Bundy et al., 2019). This sex difference in 5XFAD pathology is likely partly artifactual, since the Thy1 promotor utilized in the 5XFAD model responds to oestrogen (Sadleir et al., 2015; Poon et al., 2023).

In addition to assessing sleep changes after DPH treatment, we assessed sleep differences in our 5XFAD and WT animals. Sleep disturbances in 6– 9-month 5XFAD mice are well-characterized in many studies as decreased slow-wave sleep (SWS), shorter sleep bouts, and decreased total sleep time compared to wildtype (WT) controls (Zhang et al., 2025; Drew et al., 2023; Duncan et al., 2019; Mander, 2020; Sethi et al., 2015). These sleep deficits mirror sleep disturbances reported in human patients with Alzheimer’s disease, dementia, and cognitive impairment, supporting the use of 5XFAD as a useful Alzheimer’s model for sleep studies (Prinz et al., 1982; Bliwise et al., 2011). Sleep deficits in 5XFAD mice are typically characterized at 6 months of age or beyond, when robust memory impairments emerge. However, growing evidence indicates that sleep disturbances emerge earlier, and may serve as an early marker of disease progression (Ju et al., 2014; Parhizkar et al., 2025; Sethi et al. 2015). Motivated by this growing evidence, we first aim to characterize the prodromal sleep disruptions in 5XFAD mice. To characterize the prodromal sleep changes in 5XFAD mice, we evaluated sleep quality at 5-months-old, which is the typical age of onset of symptomatic phenotype in 5XFADs (Mar et al. 2024; Lin et al., 2023; Sil et al., 2022; Zhang et al., 2025).

## 2. Methods

### 2.1. Mice

5XFAD-B6SJL (RRID:MMRRC, JAX:034840) transgenic mice overexpressing three human amyloid beta precursor protein (APP) mutations– the Swedish (K670N, M671L), Florida (I716V), and London (V717I)– and two human presenilin 1 mutations (M146L and L286V) were obtained from Jackson labs (Bar Harbor, ME, USA) and subsequently maintained in-house by crossing male 5XFAD hemizygous mice to female B6SJLFi/j mice (RRID:IMSR, JAX:100012). Resulting offspring were either hemizygous for the 5XFAD mutations or WT for the 5XFAD mutations. WT littermate offspring were used as WT controls in this study. All mice were females between 4 and 5 months of age. 16 5XFAD mice and 14 WT mice were included in the analysis (Fig. 1). After a baseline sleep recording, some mice from both genotypes were dosed with DPH. The resulting sample sizes for the chronic sleep recording were 9 vehicle-treated 5XFAD, 7 vehicle-treated WT, 7 DPH-treated 5XFAD, and 7 DPH-treated WT. Mice had *ad libitum* access to food (NIH-31 rodent diet) and water and were maintained on a 12-hour light/dark cycle under constant temperature (70-74° F) and humidity (40-60%) control. Mice were singly housed in their home cages. All procedures were approved and performed in compliance with the NIEHS/NIH Humane Care and Use of Animals in Research protocols (NL,2021-0029).

**Figure 1.**
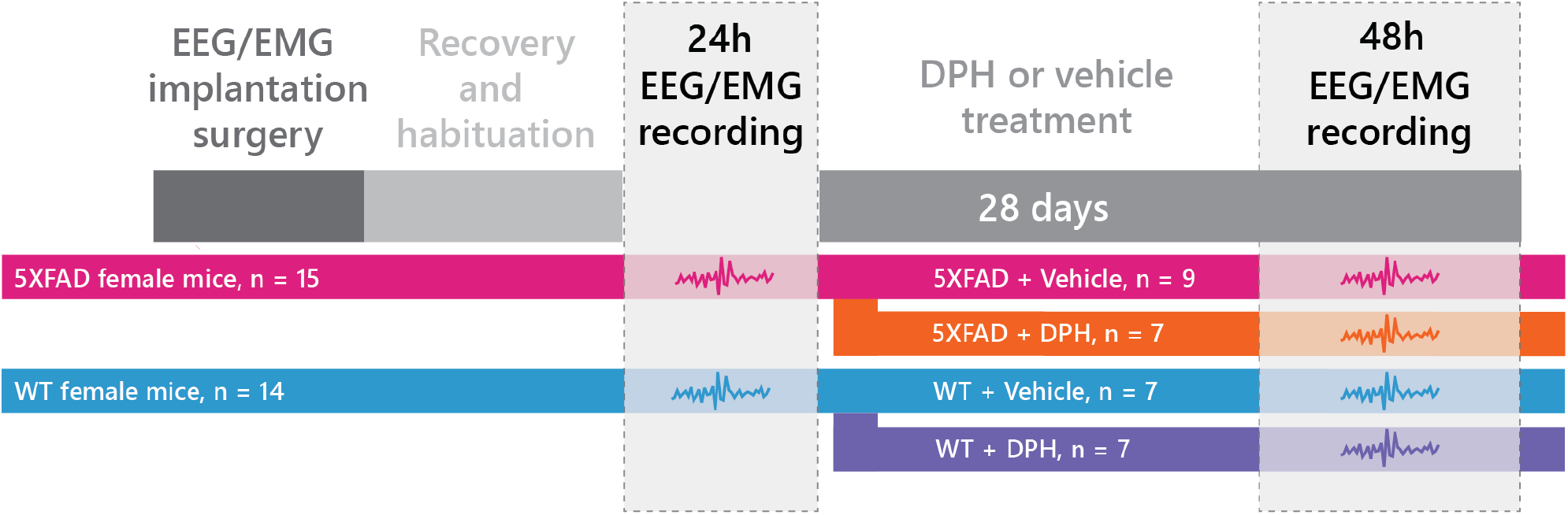
experimental design. Five-month-old female 5XFAD and WT mice were implanted with wireless EEG/EMG telemetry devices and allowed to recover and habituate before a 24h baseline sleep recording was collected to assess genotype differences in wake, REM, NREM1, and NREM2 sleep across the light and dark phases. Following baseline recording, mice received daily DPH or vehicle treatment at ZT0 for one month, with treatment groups consisting of 5XFAD + Vehicle, 5XFAD + DPH, WT + Vehicle, and WT + DPH. After one month of treatment, mice underwent a 48h EEG/EMG recording while dosing continued, allowing assessment of DPH effects on sleep architecture during a period corresponding to early cognitive decline and pathological progression in the 5XFAD model.

### 2.2. Experimental design

The experimental design is described in Fig. 1. First, all animals underwent baseline sleep recording to characterize the prodromal sleep changes in 5XFAD mice. During the baseline recording session, wireless telemetry implants were used to record EEG/EMG signal in 5-month-old female 5XFAD mice and WT controls for 24 hours. From the EEG/EMG signal, we assessed genotype differences in percentage of time in awake, REM, NREM1, and NREM2 during the lights-on period (ZT0-ZT12, hereafter the light phase) and the lights-off period (ZT12-ZT24, hereafter the dark phase). After the baseline recording, a chronic DPH dosing paradigm was initiated. In this paradigm, mice were dosed daily at Zeitgeber Time (ZT) 0 for one month. This dosing schedule spanned their 5^th^ month of life, an age at which the 5XFAD model experiences an onset of cognitive decline. This pivotal age was chosen for DPH administration to capture effects of the cycle of sleep deficits and pathology progression (Ju et al., 2014; Wang and Holzman, 2020). Because of the pathological progression spanning the 5^th^ and 6^th^ month in 5XFAD mice, a between-subject design was adopted, in which half the 5XFAD mice were dosed with a vehicle control. We tested the effect of DPH in this manner on both 5XFAD and WT mice—half the 5XFAD and WT mice received daily 10 mg/kg dosing, while the other half received vehicle control. Ultimately, four groups were compared (5XFAD+DPH, 5XFAD+Vehicle, WT+DPH, WT+Vehicle). To compare these four groups, a 48-hour long EEG recording was obtained in each mouse via wireless telemetry implants after one month of DPH treatment. Sleep recording started on the 30^th^ day of dosing (+/-6 days). DPH dosing continued during the recording period, to assess the effects of chronic ongoing DPH use on sleep quality. From this EEG/EMG signal, we assessed the effect of drug treatment on the architecture of awake, REM, NREM1, and NREM2. Using the EEG recordings, sleep states were annotated and analyzed.

### 2.3. Wireless In vivo Telemetric Electroencephalography

At four months of age, female 5XFAD-B6SJL were subcutaneously implanted with a dual EEG and EMG transmitter (HD-X02, Data Science Internationals (DSI) Harvard Bioscience, St Paul, MN, USA). The transmitter has a bandwidth of 0.5-80 Hz and a sampling rate of 1000 Hz. Standard aseptic surgical procedures were followed. Mice were anesthetized with 3-4% isoflurane in their home-cage, then maintained at approximately 1% isoflurane at a flow rate of 1 L/min for the duration of the procedure on a stereotaxic frame. Their shaved scalp was cleaned with povidone-iodine swab stick for antiseptic/germicide and they were given a 0.01mL subcutaneous injection of 0.5% bupivacaine prior to a midline incision (∼2-3cm) that exposed the skull and neck muscles. Blunt forceps were used to make a subcutaneous pocket on the mouse’s flank for the recording implant that was irrigated with sterile saline prior to placement. After exposing and cleaning the skull, two non-magnetic stainless steel screws (P1 Technologies, Roanoke, VA, USA), acting as cortical/theta electrodes, were implanted in the skull using bregma as a reference point at the left frontal cortex (A/P: +1.0, M/L -1.5) and right caudal cortex (A/P: -3.0, M/L +3.0). The negative EEG wire was wrapped around the left frontal cortex screw, and the positive EEG wire was wrapped around the right caudal cortex screw; then both screws were covered with dental cement. The EMG leads were embedded into the neck muscle. Long-acting buprenorphine (0.02 mL/ 10 g body weight) was injected at the end of the surgery prior to suturing the incision with absorbable sutures (over the dental cement) and clipping the mouse’s hindlimb nails to prevent them from scratching the incision. Mice were singly housed for the duration of the study after surgical EEG/EMG implantation. There was a one-week period of recovery and habituation prior to recording baseline sleep.

### 2.4. Dosing

Diphenhydramine HCl (DPH) was administered orally at a dose of 10 mg/kg using approximately 0.10g of Bio-Serv transgenic dough diet (Bio-Serv, CAT#S3472 Transgenic Dough Diet, irradiated, Flemington, NJ, USA) mixed with a 0.10 M diphenhydramine solution prepared in sodium chloride with 2% Pink Wilton candy coloring and peanut oil (P2144-250mL; PC code: 1003402768; CAS-No: 8002-03-7). Food coloring was used to ensure the DPH solution was evenly mixed throughout the dough diet. Dosing occurred at ZT0 (lights on).

### 2.5. Data analysis

Ponemah software version 5 (DSI, St Paul, MN, USA) was used to collect EEG/EMG signals from the recordings performed on digital transceiver plates (TRX-1; DSI, St Paul, MN, USA). NeuroScore software version 3.4 (DSI, St Paul, MN, USA) was used to assign sleep states based on the recorded EEG/EMG signals. We separated NREM1 and NREM2 in the analyses, rather than combining the two together as NREM. To analyze the data and create plots, JMP version 18 (Statistical Analysis Systems, Cary, NC), R version 4.4.3, and Microsoft Excel version 2508 were used. Figures were made using Adobe Illustrator version 29.0.

### 2.6. Statistics

Statistical tables for each figure are included in the supplement. Bout lengths per trial, light phase, and sleep type were analyzed with survival models. Kaplan-Meier curves were compared with a log-rank test. Because no drug intervention had occurred at time of baseline, proportional hazard models for the baseline trial were fit with the control and DPH animals together and genotype as the covariate. Hazard ratios were in reference to the WT group. Proportional hazard models for the chronic trial were stratified by genotype and fit with drug as the covariate. Hazard ratios were in reference to the control group. The distribution of the bout lengths were analyzed per trial, light phase, and sleep type with a negative binomial mixed model with a random effect for animal. Bouts that spanned a light-dark transition were removed from this analysis. Models for the baseline trial were fit with drug as the covariate. Model selection for the chronic trial began with an interaction model for bout length, fit with genotype, drug, and their interaction. If the interaction term was not significant at the p = 0.05 level, an additive model was fit with genotype and drug.

## 3. Results

### 3.1. Genotype differences in sleep at baseline recording

During baseline recordings, we investigated total duration of each sleep state during the light phase. Although mice are polyphasic, most of their sleep occurs during the light phase, which corresponds to their inactive period. During the light phase, 5XFAD mice exhibited altered NREM architecture. Compared to WT mice, NREM1 was increased in 5XFADs (p=0.005282, Fig. 2E). There were no differences in percent time awake, NREM2, and REM during the light phase, nor any sleep state during the dark phase (Fig. 2).

**Figure 2.**
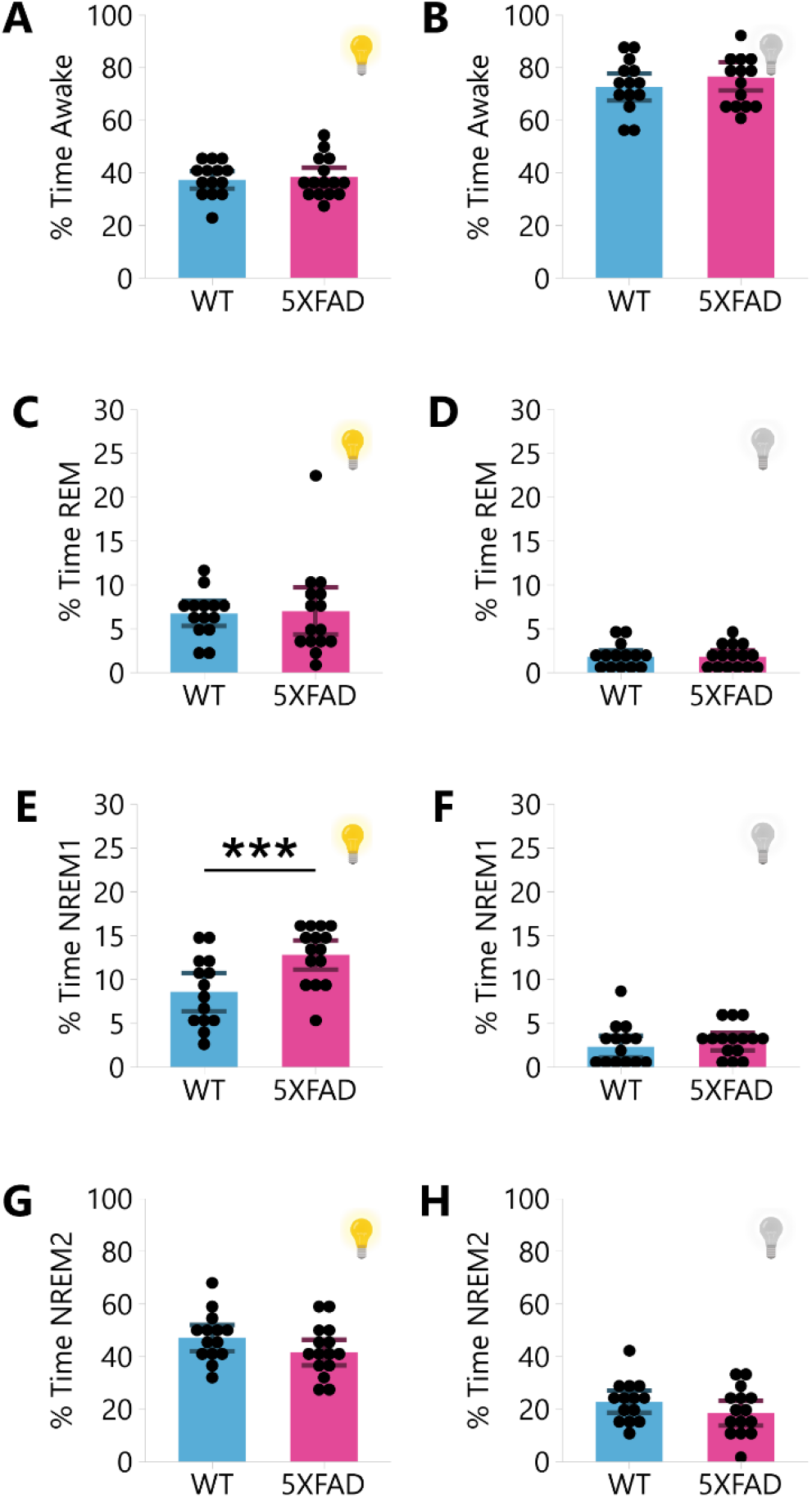
differences in sleep time in 5XFAD model. **A, C, E, G:** The mean and standard error for the percentage of time spent awake (A), in REM (C), in NREM1 (E), and in NREM2 (G) during the baseline light phase. **B, D, F, H:** The mean and standard error for the percentage of time spent awake (B), in REM (D), in NREM1 (F), and in NREM2 (H) during the baseline dark phase. Genotype differences were assessed with a t-test. ** p < 0.01.

Then, the mean bout length of awake, REM, NREM1, and NREM2 were assessed during the light phase (Fig. 3). Since mean values assume data distribution is gaussian, which bout lengths are not, the Kaplan-Meier survival probability of bouts persisting over time was also assessed; a method which avoids assumptions about data distribution (Fig. 3, S1). We found that awake bouts are not fragmented in 5XFADs (Fig. 3A-B, S1A-B). REM, however, is fragmented: the Kaplan-Meier survival probabilities show that REM sleep terminates faster for 5XFADs than WTs (p<0.0005, Fig. 3D). This REM fragmentation is confirmed using the mean bout lengths (p=0.013103, Fig. 3C). NREM1 bouts terminate faster for 5XFADs than WTs (p=0.002, Fig. 3F), though there are no differences in the NREM1 mean bout length (p=0.2461, Fig. 3E), which potentially suggests NREM1 fragmentation. NREM2 terminates faster in 5XFADs (p<0.0005, Fig. 3H) according to the Kaplan-Meier survival probabilities which indicates NREM2 fragmentation. The mean bout lengths confirm this trend (p=0.057945, Fig. 3G). These patterns of REM and NREM2 fragmentation in 5XFAD mice persist during the dark phase, where mice sleep considerably less, indicating a disruption to sleep consolidation in this genotype that is not limited to a particular time of day (REM p<0.0005, S1D, p=0.04684, S1C; NREM2 p<0.0005, S1H).

**Figure 3.**
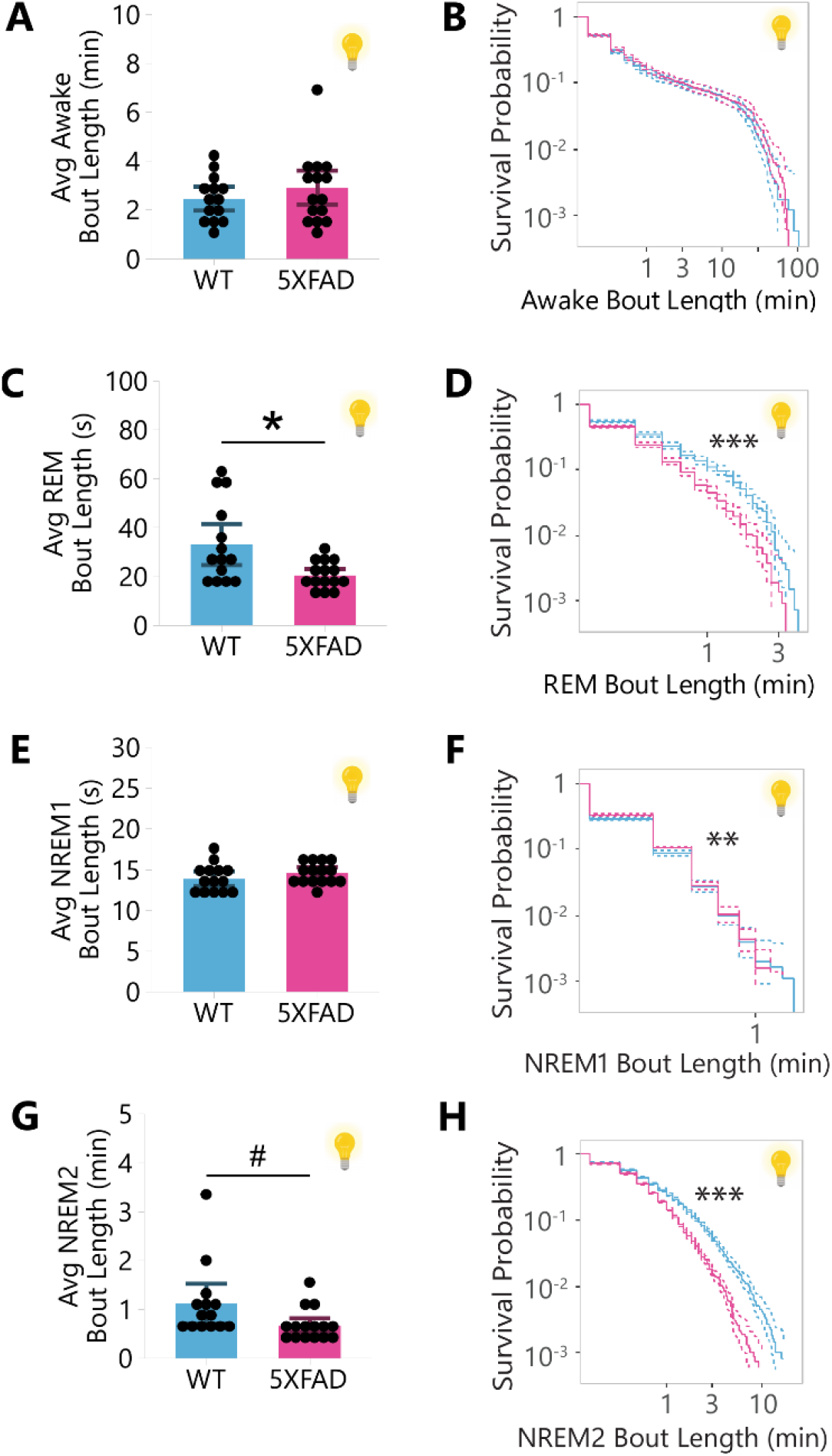
sleep fragmentation in 5XFAD model. A, C, E, G: The mean and standard error for the average bout length of awake (A), in REM (C), NREM1 (E), and NREM2 (G) in the baseline light phase. B, D, F, H: The Kaplan-Meier survival curve and 95% confidence interval for the probability that a bout of awake (B), REM (D), NREM1 (F), and NREM2 (H) will continue over time during the baseline light phase. Genotype differences were assessed with a logrank test. # p < 0.1, * p < 0.05, ** p < 0.01, *** p < 0.001.

### 3.2. DPH reduced total sleep time and NREM2 sleep during the light phase

After one month of sustained DPH treatment, sleep was recorded again as dosing continued to assess the effect of chronic ongoing DPH treatment in both 5XFAD animals and WT (Fig. 1). During this recording, we first broadly investigated the effect of sustained DPH dosing on total duration of awake, REM, NREM1, and NREM2 during the light phase (Fig. 4). We found that DPH reduced total sleep time (p_Drug_ = 0.037046, Fig. 4A). DPH also diminished time in NREM2 sleep (p_Drug_ = 0.029858, Fig. 4D). Compared to the genotype differences we observe in (Fig. 2G), DPH treatment failed to rescue the sleep deficits in 5XFAD mice; to the contrary, we saw a decrease in NREM2 sleep in both genotypes.

**Figure 4.**
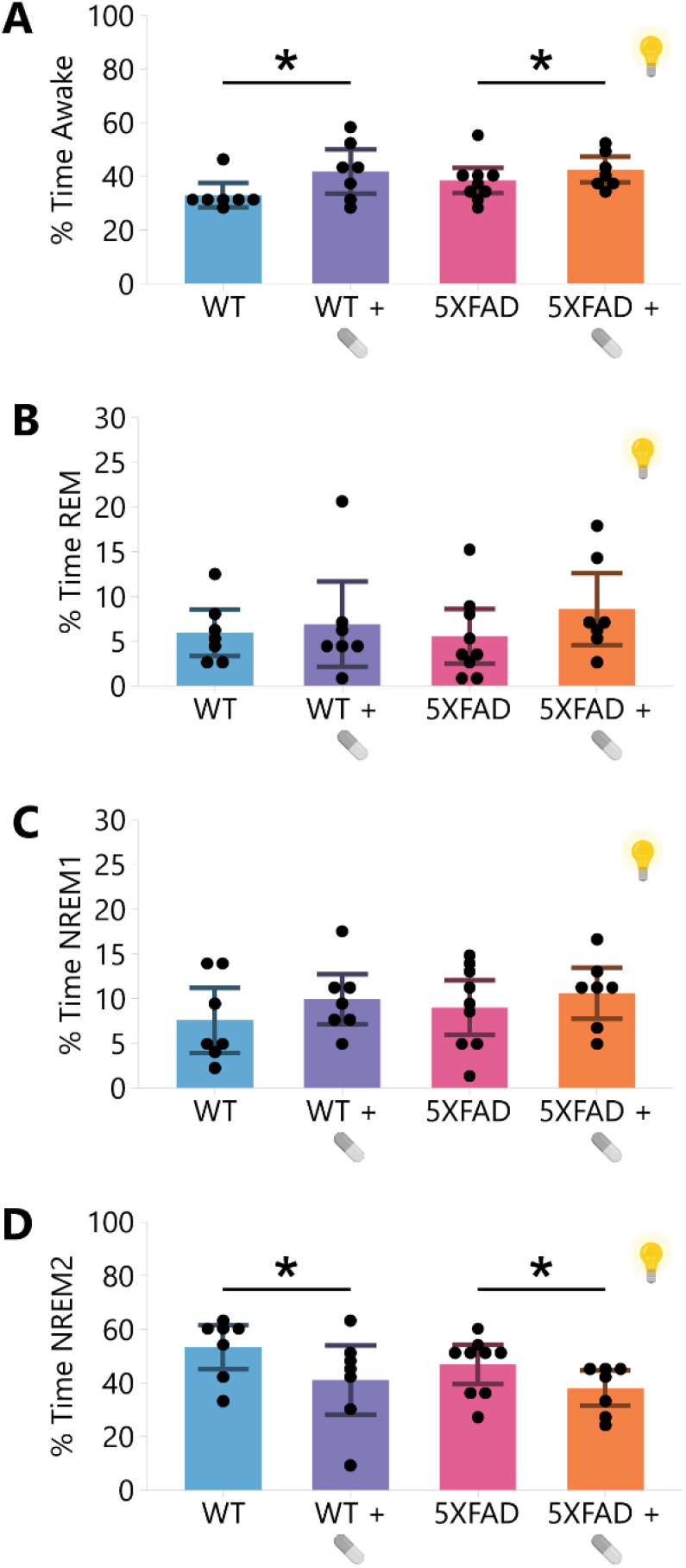
DPH reduced total sleep time and NREM2 sleep during the light phase. The mean and standard error for the percentage of time spent awake (**A**), in REM (**B**), NREM1 (**C**), and NREM2 (**D**) in the chronic light phase. The reported p-value reflects DPH from a two-factor additive ANOVA with covariates Genotype and DPH. * p < 0.05.

### 3.3. DPH increased NREM1 and REM sleep during the dark phase

We then investigated the effect of DPH on total duration of each sleep state during the dark phase, to examine whether DPH disrupted diurnal patterns of sleep. We found that DPH increased the percentage of time in NREM1 (p_Drug_ = 0.010982, Fig. 5C) and REM (p_Drug_ = 0.036471, Fig. 5B) during the dark phase in both genotypes. Together, this data indicates that DPH may have some sedating effects during the dark phase, as drug treatment promotes several sleep states during this period. However, percent time in NREM2 was not promoted by DPH (p_Drug_ = 0.807584, Fig. 5D). Therefore, the sedating effects of DPH during the dark phase do not make up for NREM2 deficits from the light phase. Moreover, since NREM2 is the most prevalent sleep state, the overall sleep/wake time was not different between DPH-treated and vehicle-treated mice (p_Drug_ = 0.265034, Fig. 5A).

**Figure 5.**
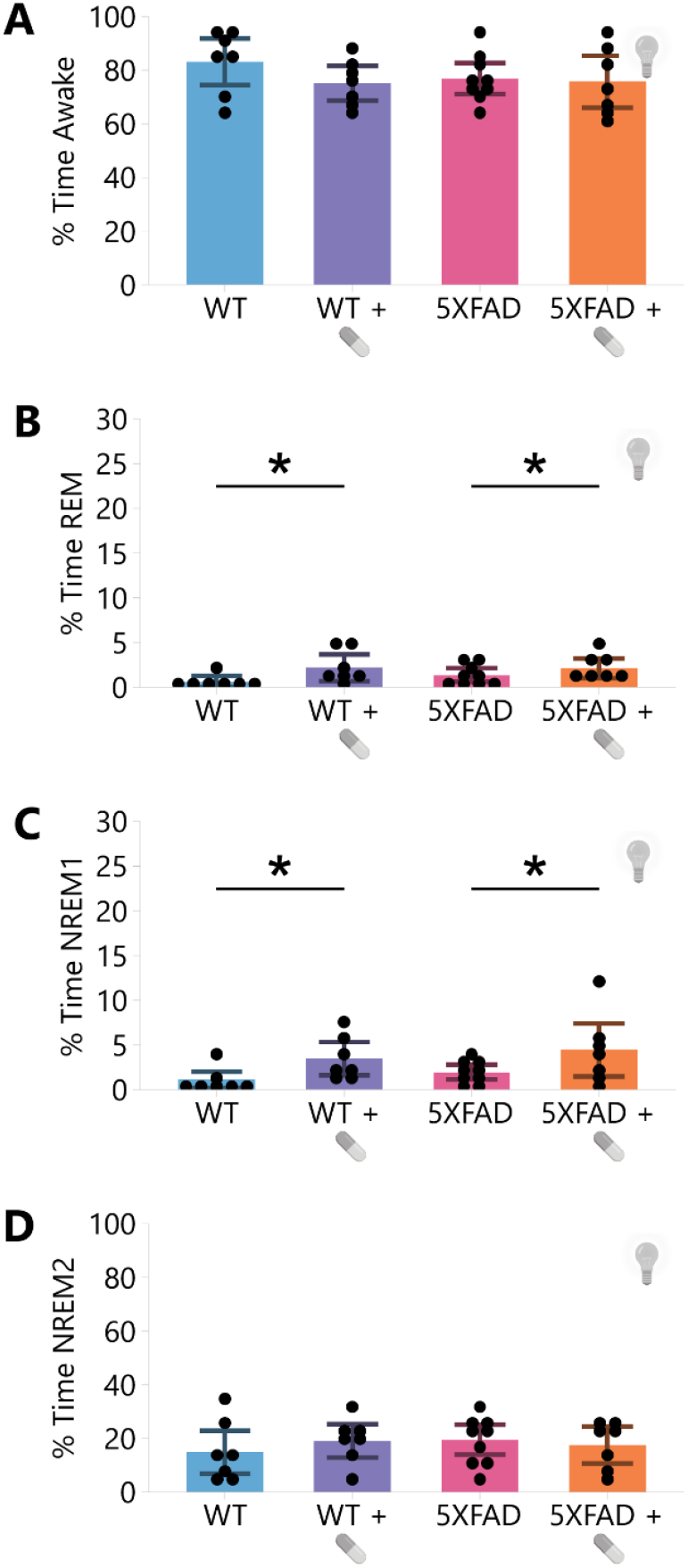
DPH increased NREM1 and REM sleep during the dark phase. The mean and standard error for the percentage of time spent awake (**A**), in REM (**B**), NREM1 (**C**), and NREM2 (**D**) in the chronic dark phase. The reported p-value reflects DPH from a two-factor additive ANOVA with covariates Genotype and DPH. * p < 0.05.

### 3.4. DPH disrupted light/dark distribution of sleep

Given that DPH increased NREM1 and REM during the dark phase, we then investigated whether DPH shifted the light/dark distribution of these elevated sleep states over 24hrs (Fig. 6A). We found that DPH pathologically disrupted the light/dark distribution in several ways, first by shifting sleep towards the dark phase rather than the light phase (p_Drug_ = 0.017862, Fig. 6B). This disruption seems to be driven by NREM1 and REM; we found that DPH-treatment shifted the distribution of REM (p_Drug_ = 0.027234, Fig. 6C) and NREM1 (p_Drug_ = 0.018978, Fig. 6D) towards the dark phase rather than the light phase. However, there was no shift in light/dark distribution for NREM2 (p_Drug_ = 0.13657, Fig. 6E). As a caveat, variability in REM sleep is intrinsically high due to relatively low prevalence of this sleep state.

**Figure 6.**
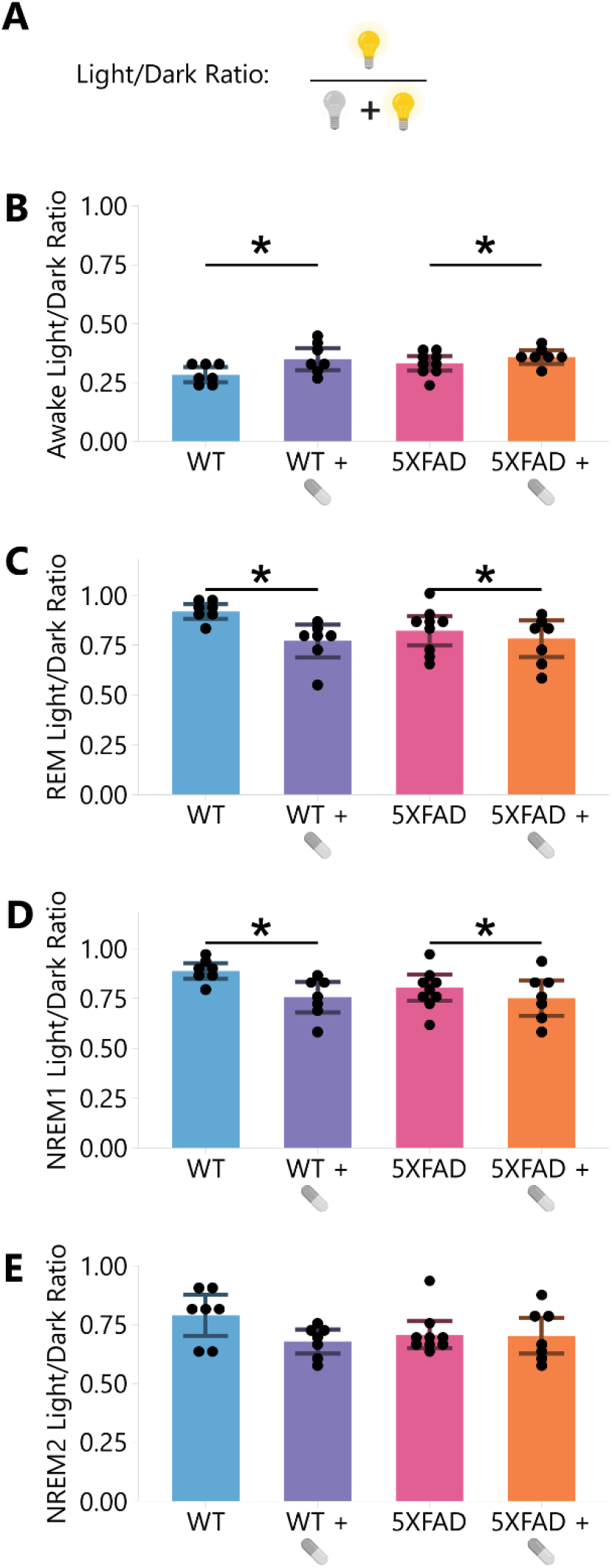
DPH disrupted light/dark distribution of sleep. **A:** Light/dark ratio is calculated as the time of a sleep phase during the light phase divided by the total time of the sleep phase over 24 hours. **B-E:** The mean and 95% confidence interval for the light/dark ratio of percent time in Awake (B), REM (C), NREM1 (D), and NREM2 (E) during the chronic phase. The reported p-value reflects DPH from a two-factor additive ANOVA with covariates Genotype and DPH. * p < 0.05.

### 3.5. DPH fragmented NREM2 sleep

We qualitatively examined NREM2 microarchitecture in representative mice (Fig. 7). One mouse from each group is illustrated; mice were chosen on the basis that their average NREM2 bout length was closest to the group average, and the representative mice are colored in grey in Fig. 8C. The NREM2 dynamics of each representative mouse are shown in Fig. 7A from ZT0 to ZT12 in a typical 12-hour light phase. Using these representative mice, we qualitatively confirmed that NREM2 appears more fragmented in the DPH-treated mice than the vehicle-treated controls. Expanding a representative 8-minute period near ZT6, we again qualitatively confirmed that NREM2 appears more fragmented in DPH-treated animals, in both genotypes (Fig. 7B-E). This fragmentation may be amplified in the 5XFAD genotype, as 5XFAD mice seem to have the poorest maintenance of NREM2 (Fig. 7E). Moreover, interruptions to NREM2 were frequently transitioned to NREM1, the “lighter” slow-wave sleep state. Together, these data indicate that DPH impaired the ability of mice to maintain NREM2 sleep, and DPH increased this impairment.

**Figure 7.**
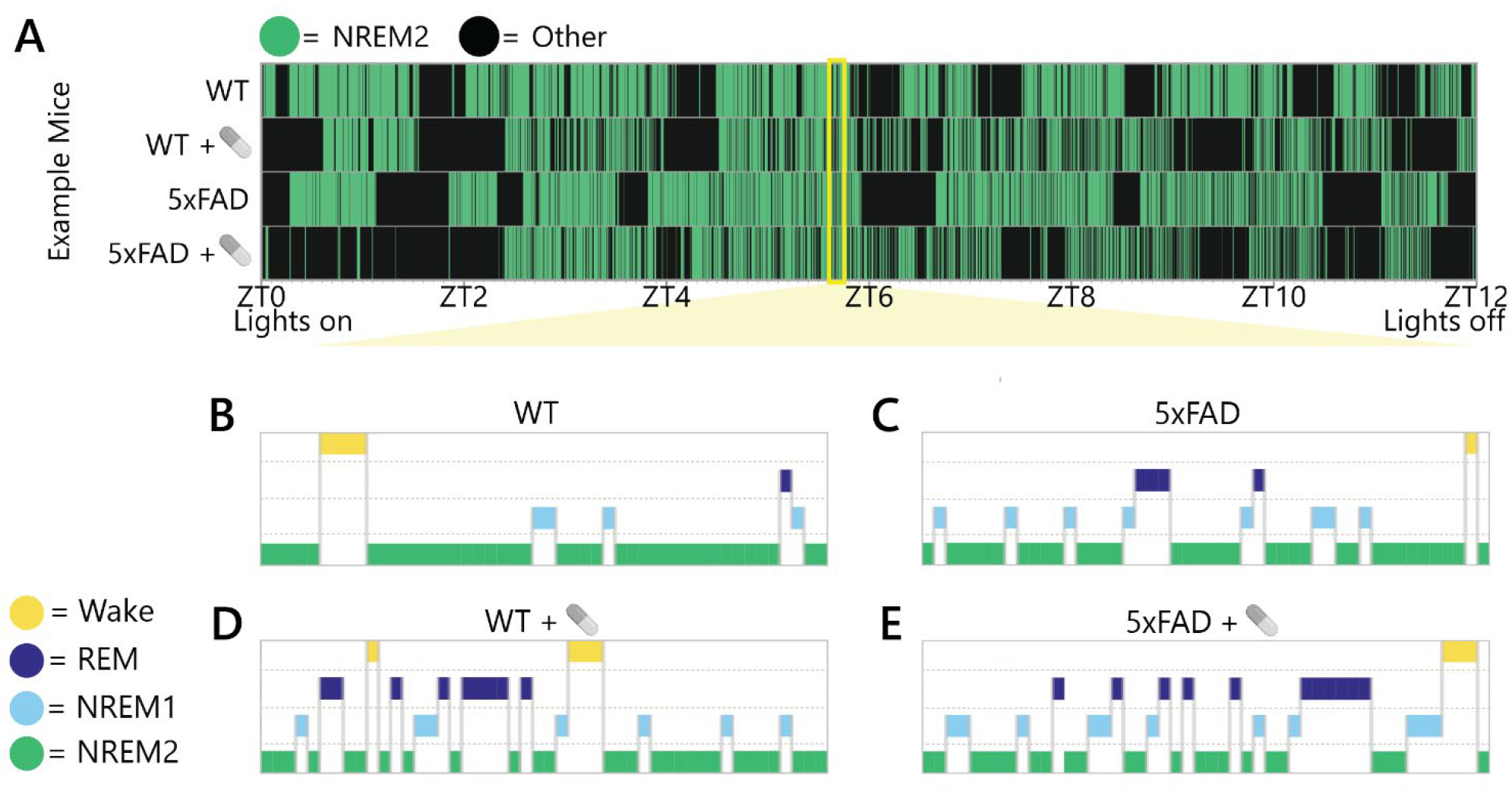
DPH qualitatively disrupted NREM2 architecture in example mice. **A:** Representative 12-hour light phase during the chronic recording displaying NREM2 compared to any other sleep phase for four example mice. **B-E:** Hypnograms during a representative 8-minute period during the light phase for the same four example mice shown in (A).

**Figure 8.**
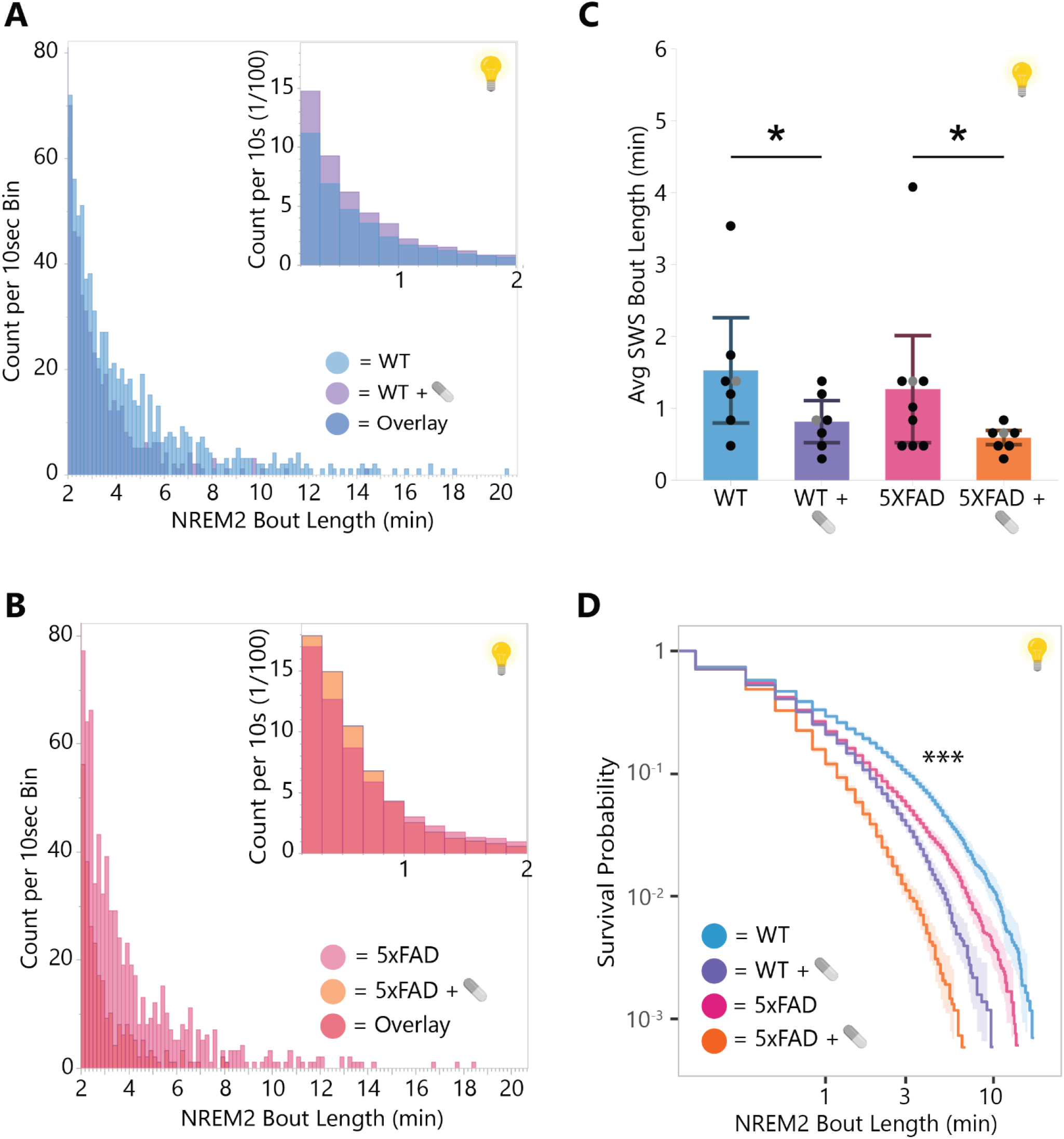
DPH fragmented NREM2 sleep. **A:** Histogram of average count of NREM2 bouts in WT mice treated with vehicle versus DPH. Inlay shows histogram for bouts less than two minutes and larger graph shows bouts more than two minutes. **B:** Histogram of average count of NREM2 bouts in 5XFAD mice treated with vehicle versus DPH. Inlay shows histogram for bouts less than two minutes and larger graph shows bouts more than two minutes. **C:** The mean and standard error for the average bout length of NREM2 in the chronic light phase. The reported p-value reflects DPH from a two-factor additive ANOVA with covariates Genotype and DPH. **D:** The Kaplan-Meier survival curve and 95% confidence interval for the probability that a bout of NREM2 will continue over time during the chronic light phase. Genotype and DPH differences were assessed with a logrank test. * p < 0.05, ** p < 0.01, *** p < 0.001.

After investigating fragmentation in each treatment group qualitatively, we quantitatively assessed sleep fragmentation after DPH treatment (Fig. 8, S2-5). Particularly, NREM2 was fragmented; DPH decreased average bout length of NREM2 for both genotypes by 42.9% (Neg. Bin. p_Drug_ = 0.004) from 77.5 seconds in WT controls to 44.3 seconds in WT DPH and from 60.3 seconds in 5XFAD controls to 34.5 seconds in 5XFAD DPH. Although the genotype averages were different, there was no statistically significant genotype effect (Neg. Bin. p_Genotype_ = 0.191) (Fig. 8C). DPH impaired the ability for mice to stay in NREM2 for a long time; bout lengths above 10 minutes of NREM2 were relatively common for vehicle treated mice (WT Control: 1.32%, 5XFAD Control: 2.10%), but virtually absent for DPH-treated mice (WT DPH: 0.08%, 5XFAD DPH: 0.00%) of both genotypes (Fig. 8A-B). Moreover, DPH increased the frequency of very short bouts (less than 10s) in both genotypes (WT Control: 24.1%, 5XFAD Control: 23.0%, WT DPH: 30.2%, 5XFAD DPH: 28.0%) (Fig. 8A-B, inset). We also found that DPH fragmented each other sleep state in both 5XFADs and WTs according to Kaplan-Meier survival curve analysis (S2B, D, F) and hazard ratios (S4, S5), though these effects were not observed in ANOVAs of mean bout lengths (S2A, C, E).

## 4. Discussion

Overall, our findings show that DPH impairs sleep architecture in wild-type controls and worsens sleep disruptions already present in the 5XFAD mice. DPH causes a pathological reduction in NREM2 sleep (Fig. 4D) and disruption of light/dark distribution (Fig. 5, Fig. 6). We observed a remarkable similarity between the 5XFAD sleep deficits and effects of DPH – particularly in sleep fragmentation (S2, S3, S4), especially NREM2 (Fig. 7, Fig. 8). These findings add to the growing evidence that anticholinergics should not be used as sedatives and sleep-aids in either healthy subjects or patients with Alzheimer’s disease (Kan et al., 2025; Gray et al., 2015; Risacher et al., 2016; Sateia et al., 2017). These findings contradict the traditional understanding of DPH as a sleep aid and sedative. Rather than increasing sleep time, we find that DPH treatment in a chronic setting decreased total sleep time, which may exacerbate existing sleep deficits in the 5XFAD model.

To our knowledge, this study is the longest DPH dosing schedule conducted on rodent models for Alzheimer disease; most studies only investigate the acute effects of DPH treatment (Wang et al., 2015; Monti et al., 1986; Mochizuki, 2022). Our sleep data is particularly rich because of our use of wireless EEG/EMG devices, with which we recorded continuously for lengthy periods, and we employed algorithms which can delineate light (NREM1) from deep (NREM2) slow-wave sleep and utilized survival curves to gain deeper insight into the bout architecture.

Since the FDA approved the marketing of diphenhydramine as an over-the-counter sleep aid in 1982 (1), DPH has remained high in popularity for its sedative properties, its efficacy for promoting sleep often goes unquestioned. However, our results contradict previous evidence and canonical wisdom, finding that DPH treatment not only shows no improvement in sleep for rodents, but has harmful effects. The commonly held belief in its efficacy is based on evidence that DPH promoted slow-wave sleep duration (Monti et al., 1986; Monti et al., 1990), but this evidence neglects to consider which depth of slow-wave sleep is promoted by DPH. We find that in mice, DPH causes an inability to maintain the higher delta power state of NREM2, which is the most restorative sleep state (Himali et al., 2023; Zheng et al., 2024; Lee et al., 2020; Nyaaba et al., 2025); meanwhile, the drug promotes the transitional NREM1 sleep state which is thought to be less restorative. These effects could be a difference in the immediate acute effects of dosing naive subjects, compared to our study, which assesses the lasting effects of sustained dosing. So overall, our findings suggest that the original evidence justifying the use of DPH as a sleep aid failed to account for a reduction in the restorative deep sleep that we observed through a more detailed analysis of sleep architecture in the present study and a longer dosing schedule.

Moreover, our results may contribute to the recently emerging link between DPH and dementia (Su et al., 2024; Kan et al., 2025; Gray et al., 2015; Risacher et al., 2016). Based on our findings, one mechanism by which DPH increases dementia risk could be by reducing sleep quality. It is well established that dementia is accelerated by poor sleep (Shi et al., 2018; Lew et al., 2021), and our novel contribution finds that DPH fragments sleep and reduces deeper slow wave sleep states in favor of shorter ones; all these sleep deficits are risk factors for dementia. To our knowledge, this is the first study investigating a biological mechanism by which DPH leads to dementia.

We also contribute a novel analysis of sleep deficits in 5XFAD animals at 5 months old. Fragmentation of REM and NREM2 bouts are a novel finding; previous characterizations have only found reductions in REM total time (Schneider et al., 2014; Drew et al., 2023) and no changes in NREM until 5.5-6 months old (Zhang et al, 2025; Drew et al. 2023; Schneider et al., 2014). We also observe a novel yet subtle change in NREM architecture. Although total time in NREM is similar between genotypes, lighter slow-wave sleep is favored over deeper, high-delta wave states in 5XFADs (Fig. 2E, G). Specifically, 5XFADs get less NREM2 sleep than controls, and more NREM1. These findings suggest this genotype experiences less restorative sleep, even before pathology develops.

Further research is needed to investigate the mechanisms underlying how DPH reduces the ability to maintain deep slow-wave sleep. Understanding this process will provide important insights into how sleep is harmed by dementia. There is a need to learn more about how AD disrupts sleep to directly treat the issue. There are reasons to believe the biology underlying sleep disruptions in the case of dementia could be very similar to those caused by DPH: in both cases, the phenotypes of sleep disruption are remarkably similar, and the cholinergic system is compromised. These similarities suggest that diphenhydramine and AD may inhibit sleep quality by similar mechanisms.

As a limitation, the implants are about 15% of the mice’s body weight, which is a heavy weight and may confound our results by disrupting mice sleep, however, our experimental design controlled for these potential effects. Another limitation to our study is the necessity of isolating mice in single housing conditions, which is known to affect sleep or cause behavioral changes which may affect sleep (Kaushal et al., 2012; Febinger et al., 2014; Egebjerg and Kornum, 2025). Single housing is required to prevent the mice from damaging the implant of a cage-mate and thereby disrupting the study. In addition, our study only utilized female mice, which show a more rapid progression of pathology in the 5xFAD model and a higher prevalence of AD in the human population. Future studies could expand our findings to male mice to determine if there are any sex differences in the response to DPH.

While we have shown novel harms associated with DPH treatment in mice, these drugs are still widely used. More research is needed to sway the canonical view that DPH has few harms for healthy adults and is effective as a sleep aid. And while warnings of cognitive impairment due to DPH use in older adults are already ubiquitous, our findings suggest that sustained DPH treatment is also harmful for sleep quality, especially in our Alzheimer mouse model. So, the long-term harms of DPH for older adults may be worse than previously known. Moreover, the efficacy of DPH as a sleep aid may be a misconception, as our findings argue that it worsens sleep quality. Our results warrant scrutiny and reconsideration of the widely accepted use of diphenhydramine as a sleep aid, and even more stringent guidelines against its use for older adults.

## CRediT authorship contribution statement

**Margaret H. Copeland:** Writing – original draft, Validation, Formal analysis, Data curation, Visualization. **Diane E. Youngstrom:** Writing – original draft, Conceptualization, Methodology, Investigation, Visualization, Project administration. **Kathryn S. Konrad:** Formal analysis, Data curation, Writing – review & editing, Visualization. **Graham H. Diering:** Conceptualization, Writing – review & editing, Visualization. **Ayland C. Letsinger:** Conceptualization, Methodology, Investigation, Writing – review & editing, Supervision, Project administration. **Leslie R. Aksu:** Methodology, Validation, Formal analysis, Data curation, Writing – review & editing, Visualization. **Jerrel L. Yakel:** Conceptualization, Resources, Writing – review & editing, Supervision, Funding acquisition. **Jesse D. Cushman:** Methodology, Resources, Writing – review & editing, Supervision, Project administration, Funding acquisition

## Acknowledgements

We thank Julia S. Lord for her expert guidance on data presentation.

## Funding/grant information

This research was supported by the Intramural Research Program of the NIH, National Institute of Environmental Health Sciences under ZIC ES103330, 1ZIAES090089, and contract GS-00F-173CA / 75N96022F00055 to Social and Scientific Systems, Inc., A DLH Holdings Corp Company. The contributions of the NIH authors are considered Works of the United States Government. The findings and conclusions presented in this paper are those of the author(s) and do not necessarily reflect the views of the NIH or the U.S. Department of Health and Human Services.

## Declaration of competing interest

The authors declare that they have no known competing financial interests or personal relationships that could have appeared to influence the work reported in this paper.

## Data availability

Data will be made available on request.

## Declaration of generative AI and AI-assisted technologies in the manuscript preparation process

During the preparation of this work, the author(s) used ChatGPT-5.5 tools to assist with literature review and with code edits to format figures. The author(s) reviewed and edited the output as needed and take full responsibility for the content of the published article.

## Notes

### Competing Interest Statement

The authors have declared no competing interest.

