## Supplementary figures and images for "Diphenhydramine Disrupts Sleep Architecture in 5XFAD Alzheimer’s Disease Model and Wild-Type Mice"

### Supplement 1

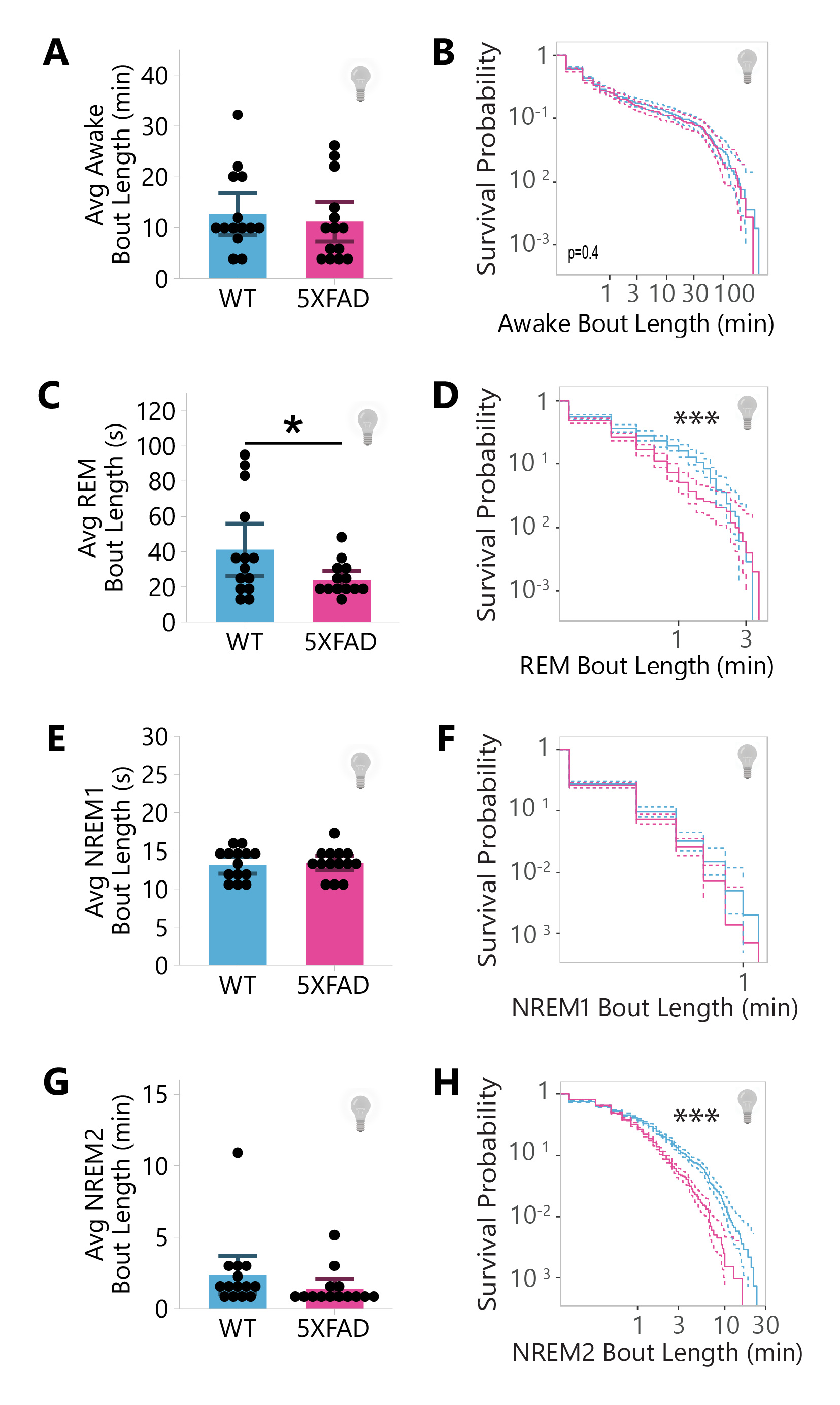

### Supplement 2

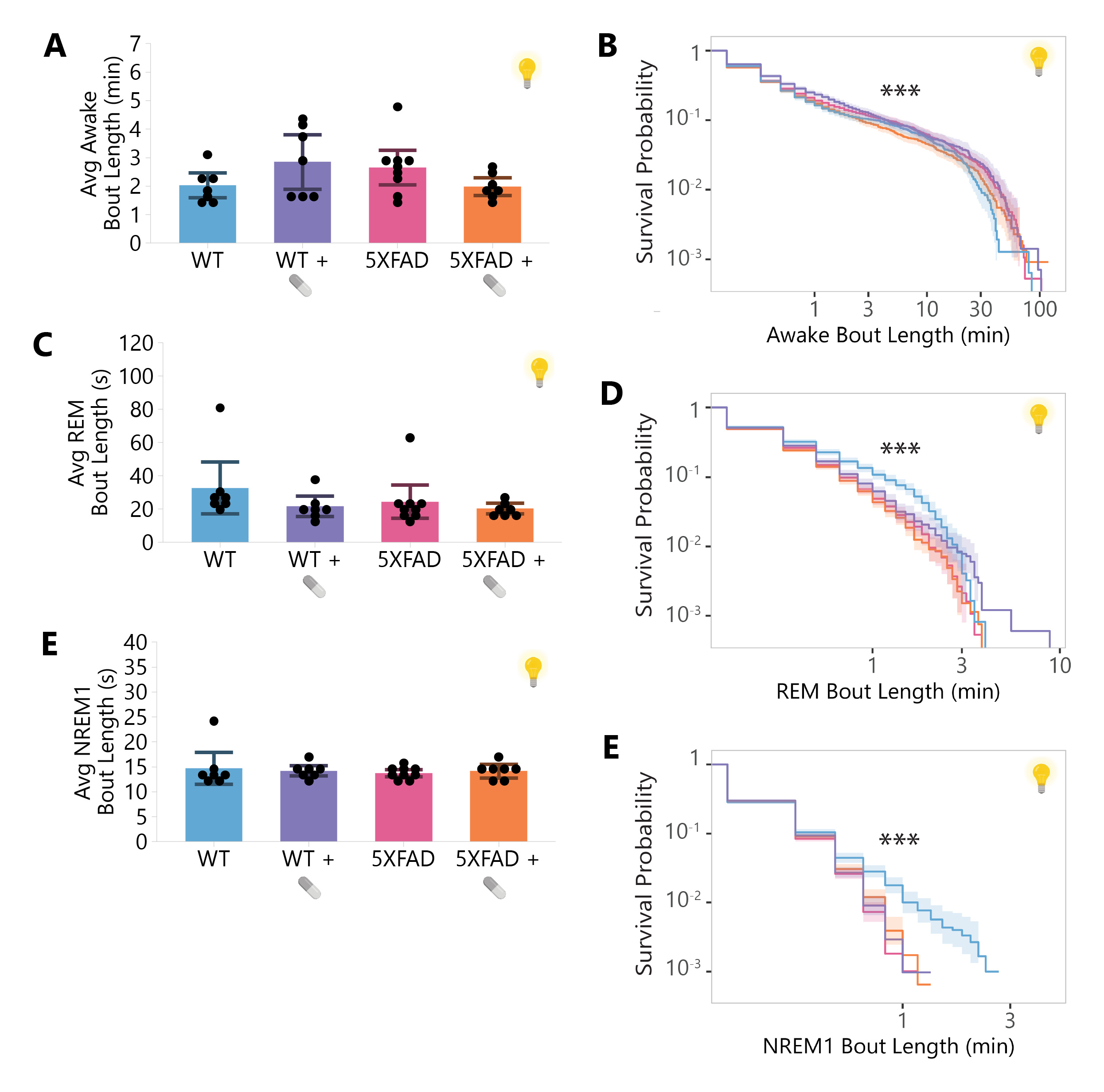

### Supplement 3

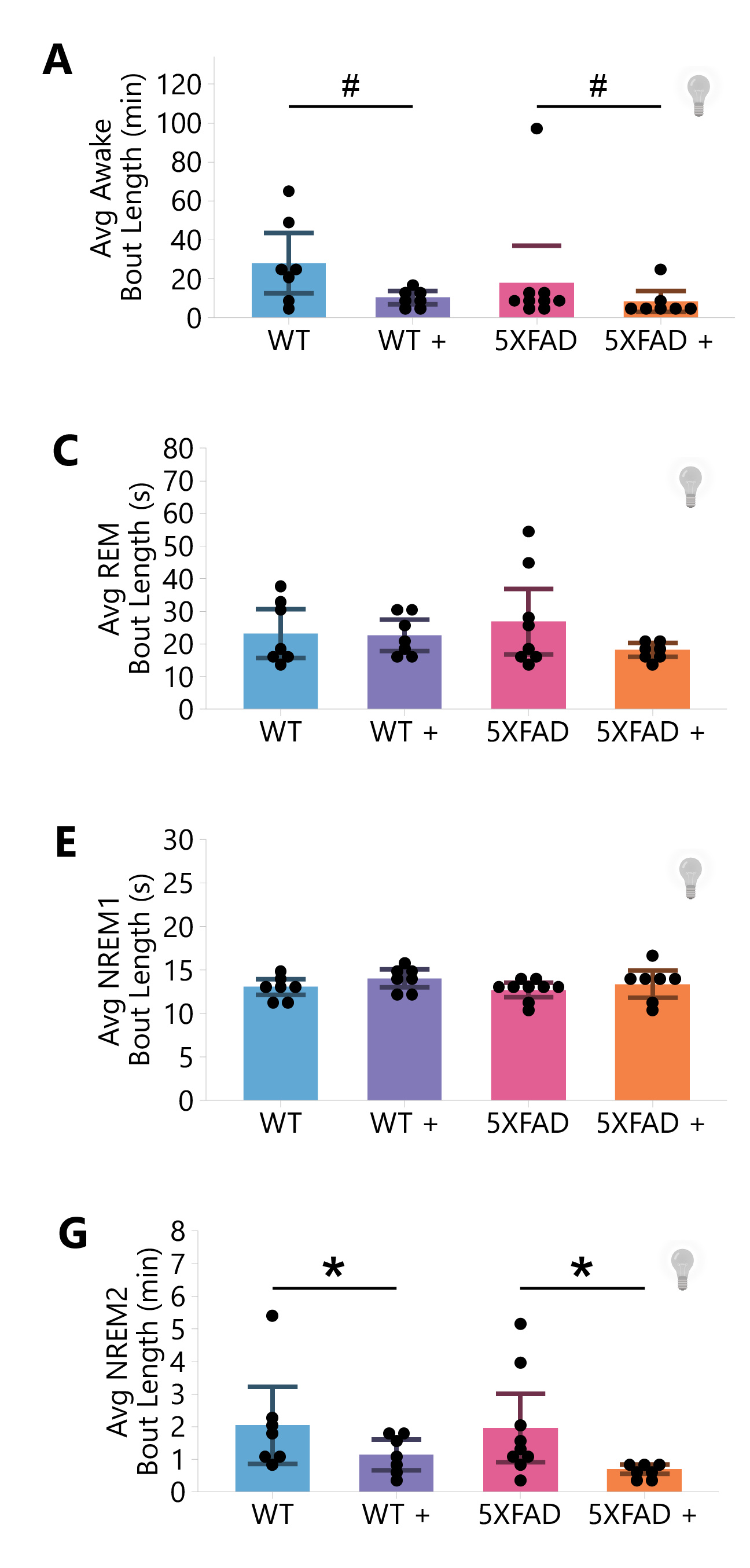

### Supplement 4

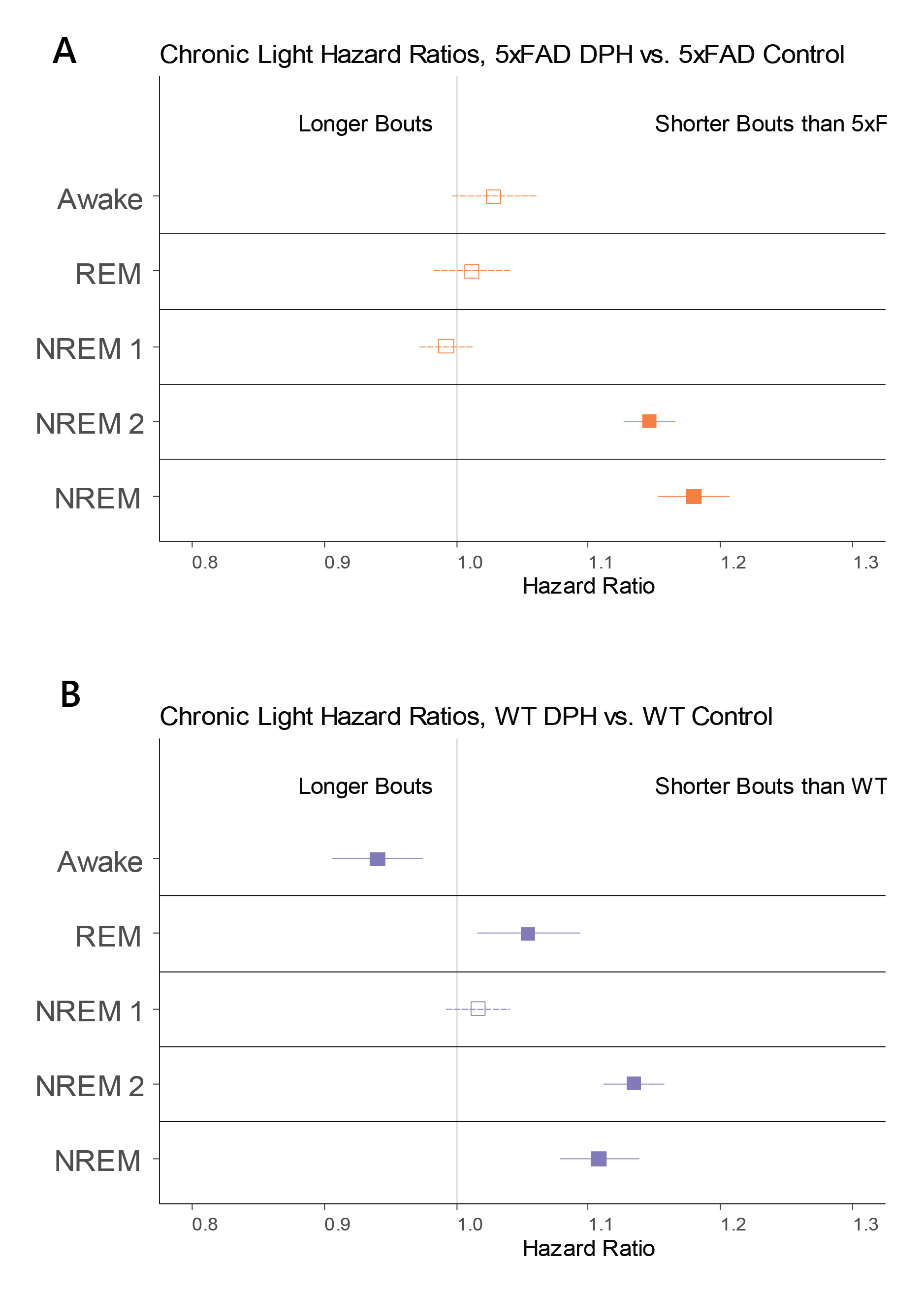

### Supplement 5

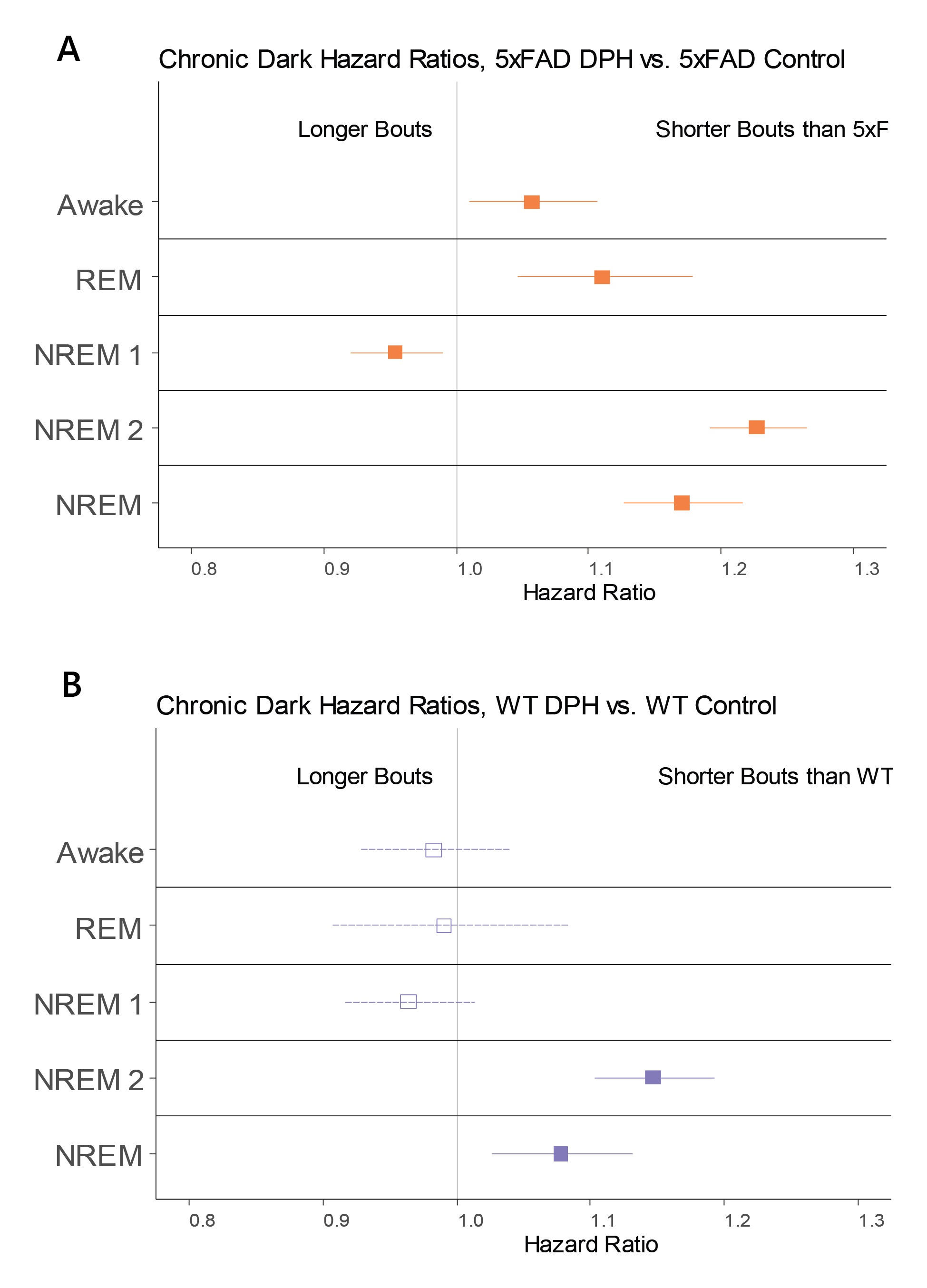

### Supplement 6

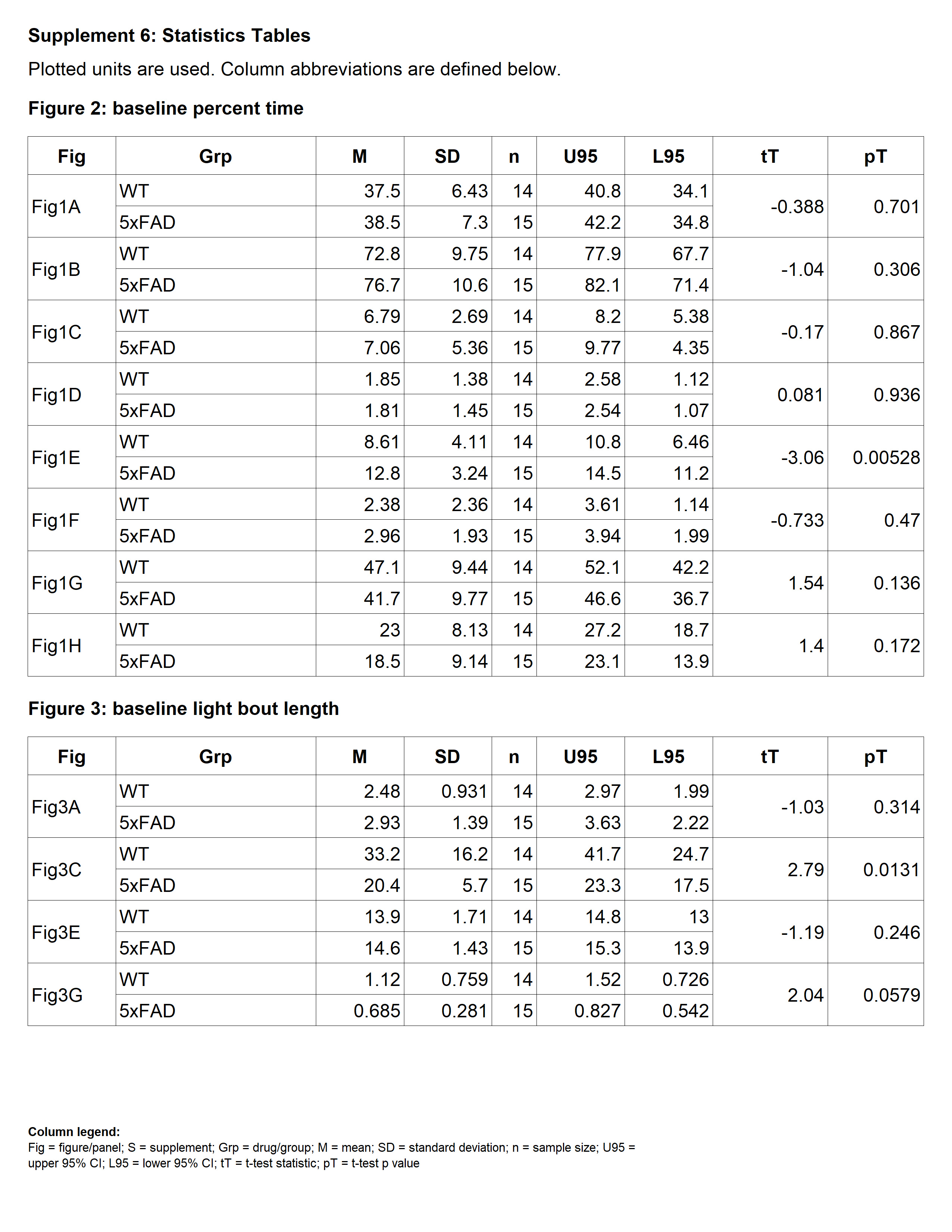

### Supplement 7

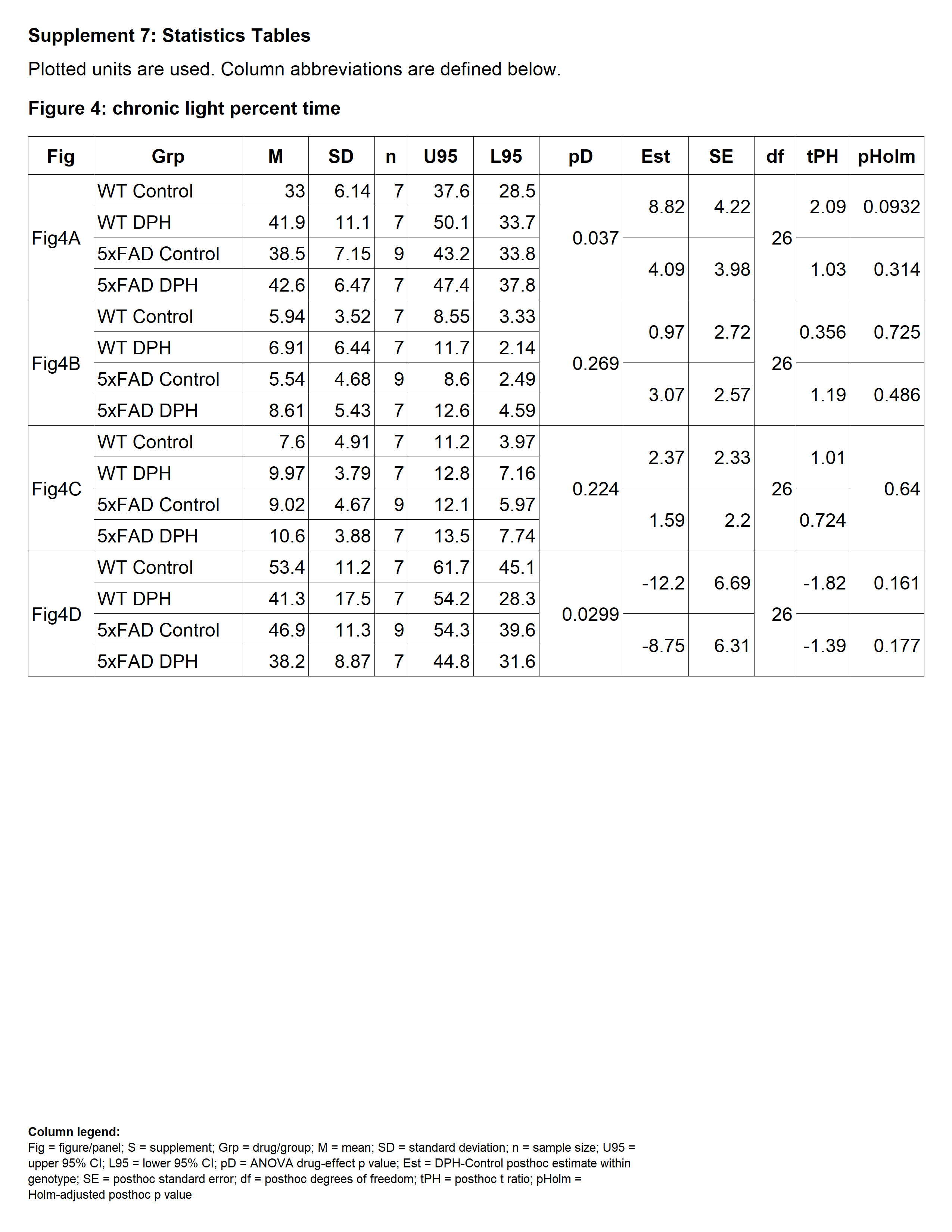

### Supplement 8

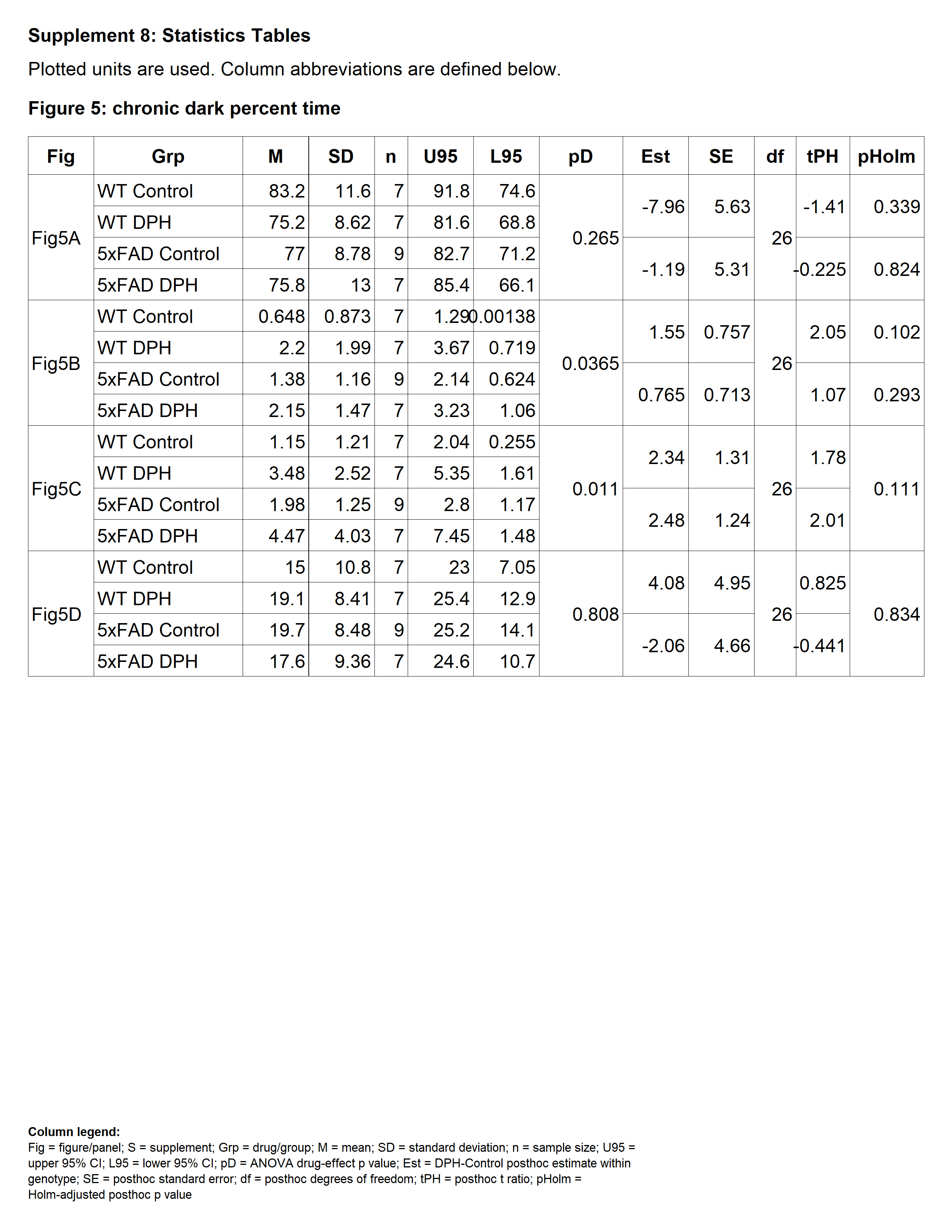

### Supplement 9

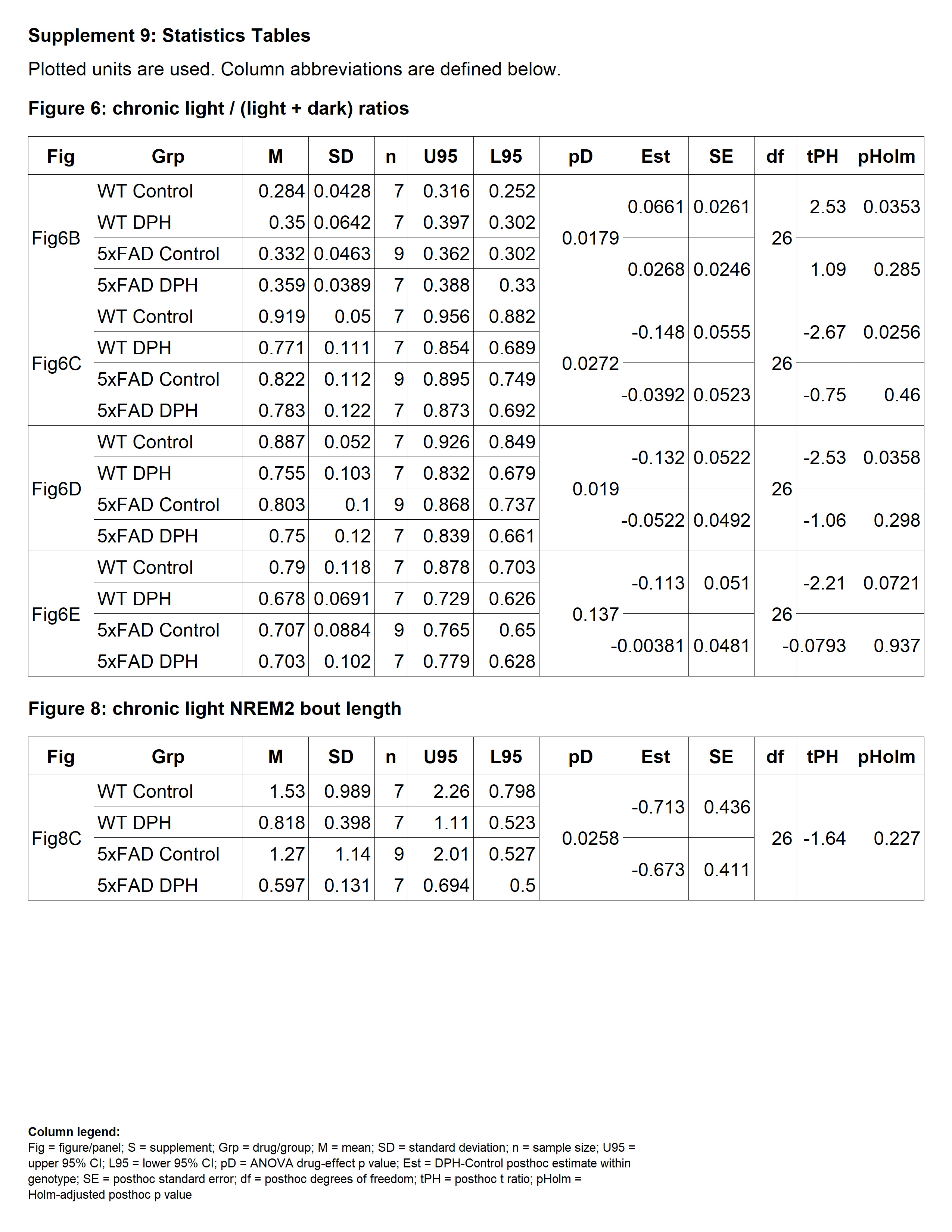

### Supplement 10

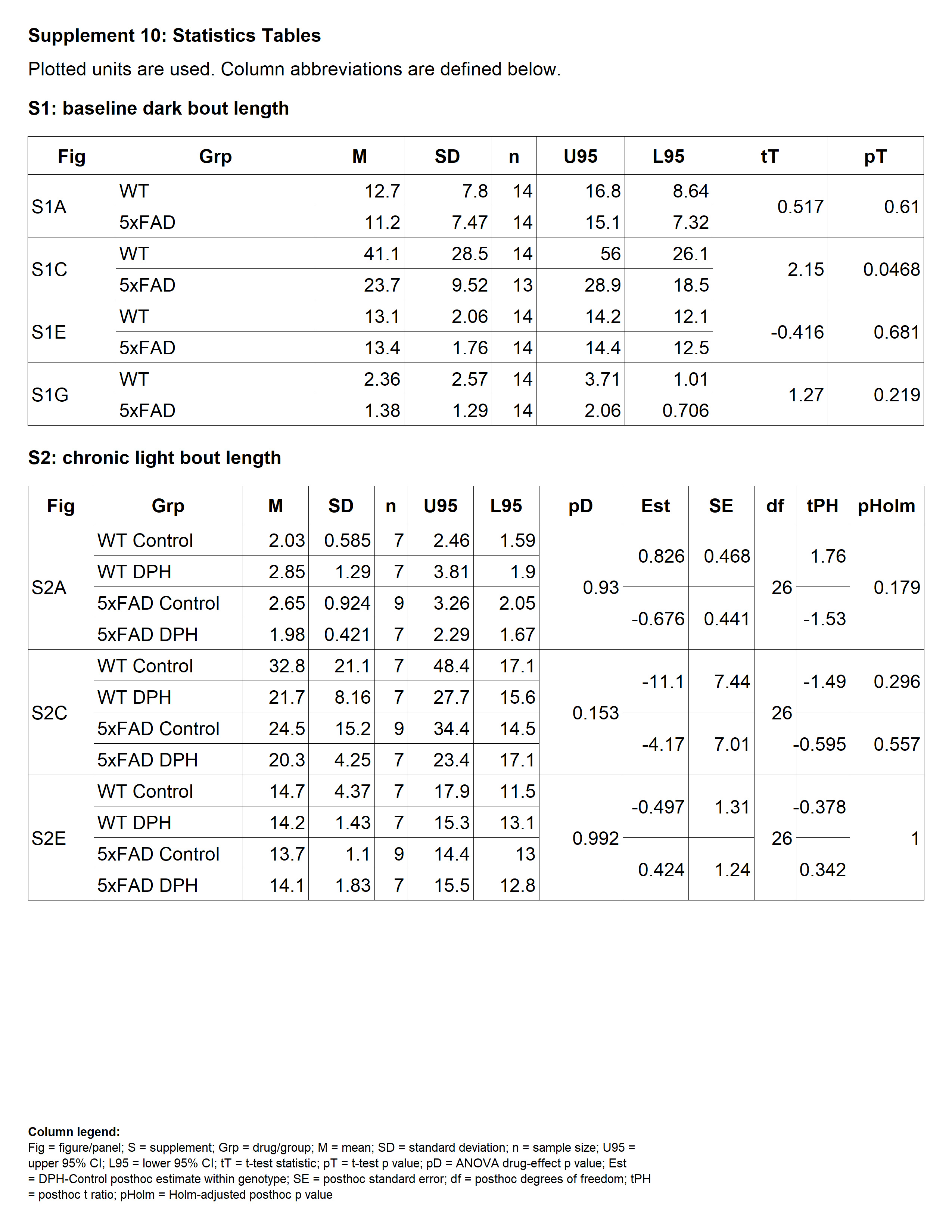

### Supplement 11

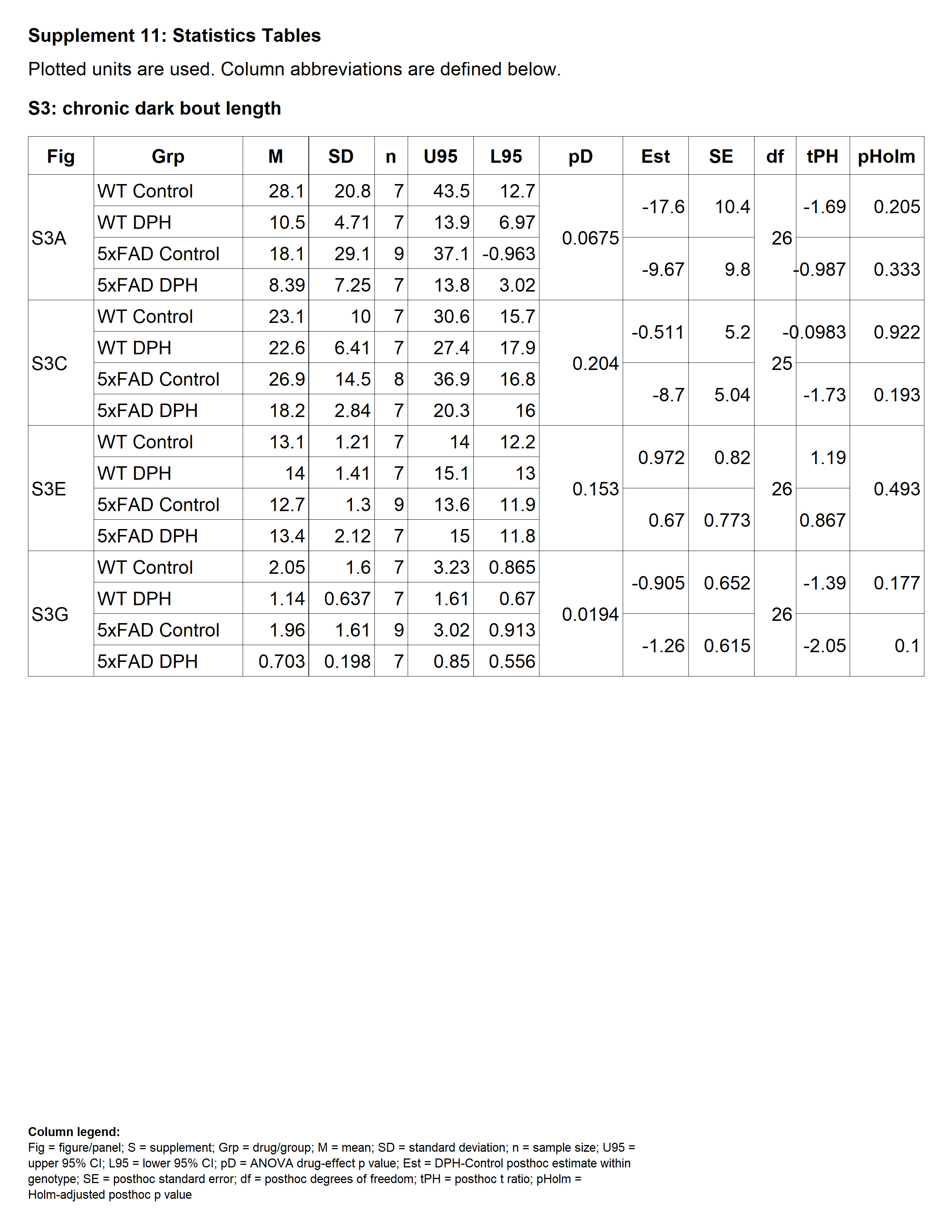

### Supplement 12

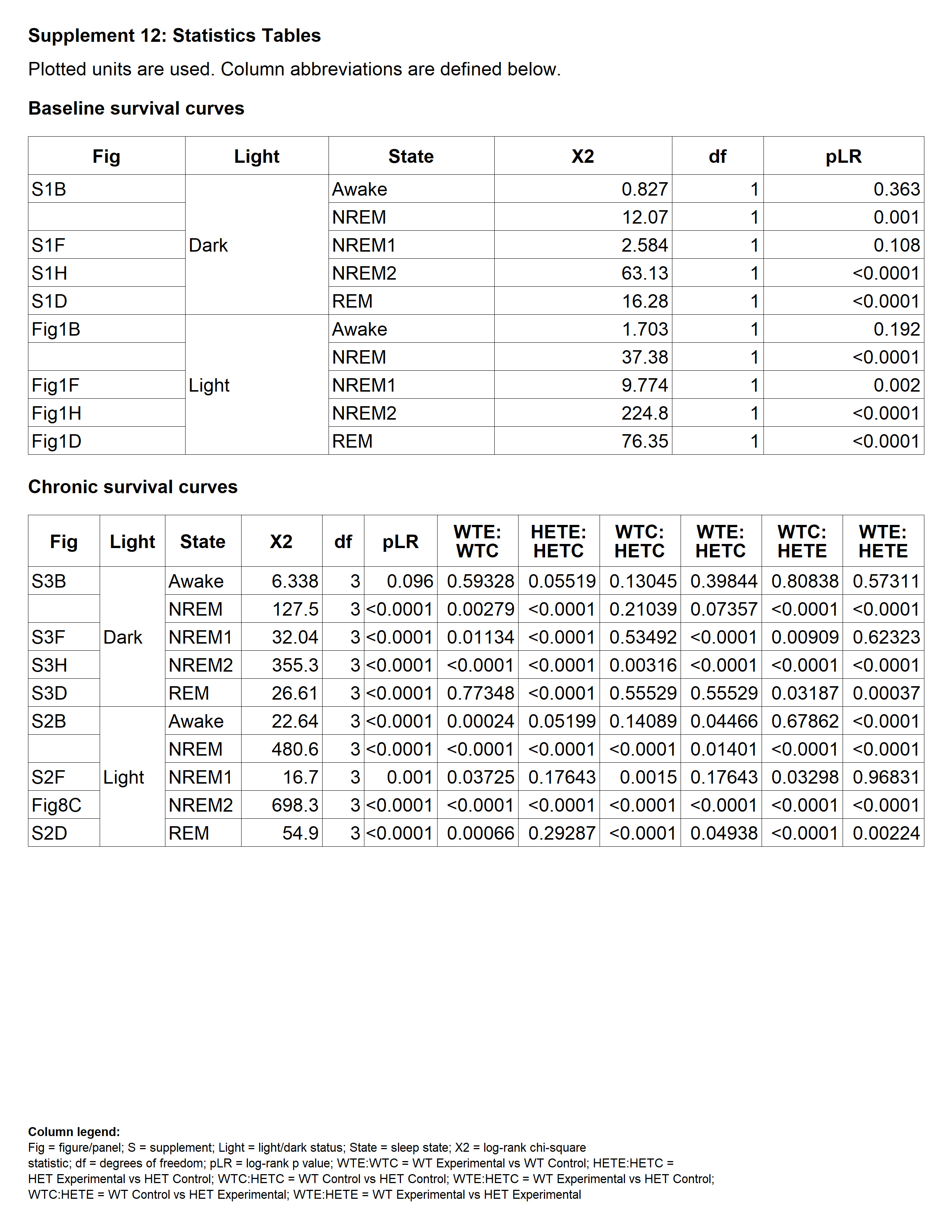
